# Acute condensin depletion causes genome decompaction without altering the level of global gene expression in *Saccharomyces cerevisiae*

**DOI:** 10.1101/195487

**Authors:** Matthew Robert Paul, Tovah Elise Markowitz, Andreas Hochwagen, Sevinç Ercan

## Abstract

Condensins are broadly conserved chromosome organizers that function in chromatin compaction and transcriptional regulation, but to what extent these two functions are linked has remained unclear. Here, we analyzed the effect of condensin inactivation on genome compaction and global gene expression in the yeast *Saccharomyces cerevisiae*. Spike-in-controlled 3C-seq analysis revealed that acute condensin inactivation leads to a global decrease in close-range chromosomal interactions as well as more specific losses of homotypic tRNA gene clustering. In addition, a condensin-rich topologically associated domain between the ribosomal DNA and the centromere on chromosome XII is lost upon condensin inactivation. Unexpectedly, these large-scale changes in chromosome architecture are not associated with global changes in transcript levels as determined by spike-in-controlled mRNA-seq analysis. Our data suggest that the global transcriptional program of *S. cerevisiae* is resistant to condensin inactivation and the associated profound changes in genome organization.

**Significance Statement:** Gene expression occurs in the context of higher-order chromatin organization, which helps compact the genome within the spatial constraints of the nucleus. To what extent higher-order chromatin compaction affects gene expression remains unknown. Here, we show that gene expression and genome compaction can be uncoupled in the single-celled model eukaryote *Saccharomyces cerevisiae*. Inactivation of the conserved condensin complex, which also organizes the human genome, leads to broad genome decompaction in this organism. Unexpectedly, this reorganization has no immediate effect on the transcriptome. These findings indicate that the global gene expression program is robust to large-scale changes in genome architecture in yeast, shedding important new light on the evolution and function of genome organization in gene regulation.

## Introduction

Organized compaction of eukaryotic genomes regulates important nuclear processes including transcription, replication, and repair (1, 2). Classic examples of nuclear organization include compartmentalization of transcriptionally active and repressed domains, as well as specialized structures as in the segregation of the ribosomal DNA (rDNA) into the nucleolus (3-5). Recent measurements of pairwise interaction frequencies between chromosomal sites revealed additional structural elements, including stable topologically associating domains (TAD) on the scale of 0.1-1Mb (6-8), but also smaller more dynamic interaction domains linked to gene regulation (9-12). Restriction of chromosomal interactions within TADs is thought to support enhancer-promoter specificity and is important for normal development (13-16).

The evolutionarily conserved condensins have emerged as major regulators of chromosomal architecture. Condensins belong to the <u>s</u>tructural <u>m</u>aintenance of <u>c</u>hromosomes (SMC) family of chromosome organizers, which also include cohesin and the SMC5/6 complex (17, 18). Like other SMC complexes, condensin forms a ring-like structure and topologically encircles DNA to mediate chromosomal looping (19, 20). Condensins are essential for chromosome condensation and segregation during cell division. In addition, condensin complexes regulate nuclear organization in interphase in a number of organisms (21-30). Interphase activities include the organization of TADs on specific chromosomes or across the genome (24, 31) and regulating interchromosomal interactions. For example, condensin complexes control X chromosome organization in *C. elegans* (24), chromosome pairing in *Drosophila* (32), and clustering of cell cycle-regulated genes in *S. pombe* (33). In *S cerevisiae*, condensin is required for tethering the centromere of chromosome XII (ChrXII) to the rDNA, rDNA compaction upon starvation, and tRNA gene clustering (34-37). Many of these specific condensin-mediated interactions were shown to have important functions in gene regulation (24, 38). It remains to be established if the general genome compaction by condensin similarly results in global effects on gene expression.

In this study, we addressed the relationship between condensin-mediated chromatin compaction and global gene expression in *S. cerevisiae* using genome-wide chromosome conformation capture (3C-seq) in wild-type, condensin-depleted and condensin-mutant cells. We find that loss of condensin function leads to a shift from shorter to longer-range chromosomal interactions, indicating that condensin helps restrict pairwise interactions to nearby loci within the genome. Our data also show a fundamental condensin-dependent organization of ChrXII, the sole rDNA-bearing chromosome in yeast. Intriguingly, these architectural changes were not associated with broad changes in global gene expression levels. These data suggest that the structural changes associated with condensin depletion are not directly coupled to transcription.

## Methods

### Cell culture and preparation of nuclei

Inducible nuclear depletion of condensin was achieved using the anchor away approach (39), specifically by tagging the condensin subunit Brn1 in the SK1 background.

Asynchronous cells were grown in 200 ml of YPD to OD1 at 30°C. Condensin depletion was induced using rapamycin at 1μg.ml^-1^ for 30 mins. Temperature-sensitive condensin mutants harbor the *ycs4-2* allele in the SK1 background (40, 41). Wild type and mutant yeast cells were grown asynchronously at permissive temperature (23°C) until OD1, then shifted to restrictive temperature (37°C), or kept at permissive temperature, for 2 hrs. For 3C-seq experiments, yeast cells were fixed for 30 mins at room temperature by adding formaldehyde to the media at 3% and quenching using 125mM glycine. Spheroplasts were prepared by digesting the cell wall with zymolyase (amsbio, UK) and then resuspending in dounce buffer (0.35M sucrose, 15mM Hepes KOH, 0.5mM EGTA, 5mM MgCl2, 10mM KCl, 0.1mM EDTA, 0.5% Triton X-100 and 0.25% NP40). Nuclei were released using a type B dounce and glass pestle on ice. Cellular debris was removed by spinning at 230g. The supernatant was spun down at 2000g to isolate the nuclei. Nuclei were stained with the vital dye Crystal violet in citric acid (Fisher Scientific, PA, USA) to allow counting on a hemocytometer as described previously (42).

### 3C-seq

For the anchor-away samples, 3C-seq was used as previously published (6). ∼100 million nuclei were permeabilized by incubating in 500μl NEB restriction enzyme buffer (New England Biolabs, MA, USA) with 0.3% SDS for 1 hr at 37°C in a thermomixer. SDS was quenched by adding Triton X-100 to 2%, and nuclei were incubated for 1 hr further at 37°C in a thermomixer. Chromatin was digested using 1200U of restriction enzyme HindIII (New England Biolabs, MA, USA) overnight at 37°C in a water bath. Following digestion, the volume was increased to a total of 8ml using 1x T4 DNA Ligase Buffer (New England Biolabs, MA, USA). Chromatin fragments were ligated with 750U of T4 DNA ligase overnight at 15°C (Thermo Fisher Scientific, MA, USA). For the temperature-sensitive samples, *in situ* 3C-seq was performed based on two protocols (6, 43). Briefly, nuclei were permeabilized by incubating in 50μl of 0.5% SDS for 10 mins at 62°C in a thermomixer. SDS was quenched by adding 170μl of 1.5% Triton X-100, and nuclei were incubated for 15 mins at 37°C in a thermomixer. 25μl of 10X NEB restriction enzyme buffer was added and chromatin was digested using 1200U of restriction enzyme HindIII overnight at 37°C in a water bath. Following digestion, the volume was increased to a total of 1.15ml using 1x T4 DNA Ligase Buffer (New England Biolabs, MA, USA). Chromatin fragments were ligated with 150U of T4 DNA ligase overnight at 15°C (Thermo Fisher Scientific, MA, USA). 3C DNA was purified using Qiagen PCR purification columns. DNA was sheared using a Bioruptor Pico (Diagenode, NJ, USA) by diluting 500ng of purified DNA in 50μl and sonicating with the following settings: 15 secs ON and 90 secs OFF for 7 cycles, with a quick vortex and spin-down after 4 cycles. Libraries were prepared from the sonicated DNA by PCR following the addition of Illumina TruSeq adapters as described previously (26). Sequencing was performed at the NYU Biology Genomics core using paired-end 50-bp sequencing on the Illumina HiSeq-2500.

### 3C-seq spike-in

To a subset of samples, we spiked in 10% *C. elegans* nuclei prior to continuing with the 3C-seq protocol above. *C. elegans* N2 embryos were washed with 500μl of chitinase buffer (110mM NaCl, 40mM KCl, 2mM CaCl_2_, 2mM MgCl_2_, 25mM HEPES-KOH pH 7.5) and digested with 5U of chitinase (Sigma-Aldrich, MO, USA). The resulting blastomeres were spun-down at 2000g for 5 mins at 4°C then resuspended in PBS with 1% formaldehyde and fixed for 30 mins while nutating. Formaldehyde was quenched with 125mM glycine. Blastomeres were washed and nuclei released in dounce buffer (see yeast nuclei preparation). Cellular debris was removed by spinning at 100g. The supernatant was spun at 2000g to isolate the nuclei.

### 3C-seq data analysis

The data statistics, such as read number and mapping efficiency, for each replicate are given in Supplemental File 1. Data were processed using previously described methods, using the Mirny lab python packages (<u>https://bitbucket.org/mirnylab/</u>) (44). Briefly,iterative mapping was used to map each side of the paired-end read individually. For the spike-in experiments, the reads that uniquely mapped to either *C. elegans* (WS220) or *S. cerevisiae* (SacCer3) were filtered and used separately for each species. Reads were then assigned to the closest restriction fragment end. Counts were binned into 10kb bins. Default filters were applied to remove outlier regions such as those with poor coverage (bottom 1% or with less than 50% of bin sequenced) or contain PCR duplicates (top 0.05%). Iterative correction was used to correct for potential biases including mappability and GC content, without pre-defining what these biases are.

### ChIP-seq and data analysis

Chromatin immunoprecipitation was performed as described (45). Samples were immunoprecipitated with 20μl anti-V5 agarose affinity gel (Sigma). ChIP and input DNA was resonicated using a Bioruptor Pico (Diagenode, NJ, USA) with the following settings: 30 secs ON and 30 secs OFF for 5 cycles. Libraries were prepared by PCR following the addition of Illumina TruSeq adapters as described previously (26). 50-bp single-end sequencing was accomplished on an Illumina HiSeq 2500 instrument at the NYU Biology Genomics core. Sequencing reads were mapped to the SK1 genome (46) using Bowtie (47). Reads that mapped to only one location without mismatches were used in further analyses. Further processing was completed using MACS-2.1.0 (<u>https://github.com/taoliu/MACS</u>) (48). Reads were extended towards 3’ ends to a final length of 200bp and probabilistically determined PCR duplications were removed. Pileups of both the input and ChIP libraries were SPMR-normalized (signal per million reads), followed by a calculation of the fold-enrichment of the ChIP data over the input data, resulting in the ChIP-coverage scores that are shown in the figures.

### mRNA-seq with S. pombe spike-in

4.5ml of OD1 *S. cerevisiae* was mixed with 500μl of *S. pombe* (972h) at OD1. To ensure that the ratio of *S. pombe* to *S. cerevisiae* was consistent between samples, genomic DNA was isolated from 10% of the mixed cells and used for qPCR. Briefly, serial dilutions of *S. pombe* and *S. cerevisiae* were amplified using both *S. pombe* and *S. cerevisiae* specific primers (Supplemental File 1) to generate standard curves, to which the amplification from experimental samples was compared. qPCR measured ratios of spike-in to experimental sample ranged between ∼10%+/-2 (values for each provided in Supplemental File 1). The cells in the remaining 90% were broken with glass beads and

RNA was purified over Qiagen RNeasy column following the recommended yeast protocol. mRNA was selected using Sera-Mag oligo-dT beads (GE Healthcare Life Sciences, Germany) from 0.5-10μg of total mRNA. cDNA was prepared based on incorporation of deoxyuridine triphosphates (dUTPs) during cDNA synthesis using a previously described protocol (49). Libraries were prepared from the cDNA by PCR following the addition of Illumina TruSeq adapters (26). Sequencing was performed at the NYU Biology Genomics core using single-end 50-bp sequencing on the Illumina HiSeq-2500.

### mRNA-seq analysis with S. pombe spike-in

*S. pombe* (ASM294v2) and *S. cerevisiae* (SacCer3) were mapped using TopHat version 2.0.11 (50) and reads mapping to both genomes were discarded from further analyses. Raw counts for each gene were determined using htseq-count version 0.6.1 (51) and normalized based on total reads. Correction of *S. cerevisiae* based on *S. pombe* spike-in was done as previously published (52). Briefly, a linear model was fitted to *S. pombe* spike-in data, comparing read counts of expressed genes (CPM >0.5) in the wild type to other experimental conditions. The coefficients from this linear model were then used to correct the *S. cerevisiae* expression values. Clustering was performed in R version 3.3.2 with hierarchical clustering using the complete linkage model. Dendrograms were cut to produce 4 clusters, as predicted by the elbow method, silhouette analysis and the Calinksi criterion.

## Results

To assess how condensin regulates chromosomal interactions in *S. cerevisiae* (yeast), we performed 3C-seq analyses, which capture the frequency of pair-wise contacts between genomic loci (6, 24). Unlike metazoans, yeast contains a single condensin complex (Fig. 1A). We acutely removed condensin from yeast nuclei using the inducible anchor-away system targeted against the kleisin subunit Brn1. In this system, proteins tagged with the FRB domain of human TOR kinase can be conditionally depleted from the nucleus by addition of rapamycin (Fig. 1B) (39). Validating this approach, the parental anchor-away strain was capable of growing on rapamycin plates whereas a strain containing FRB-tagged Brn1 was not (Fig. S1A). In an independent approach, we used a strain bearing a temperature-sensitive mutation in the Ycs4 subunit (*ycs4-2*) to conditionally inactivate condensin by shifting cells to the non-permissive temperature of 37°C (41). Sequencing showed that this allele harbors two mutations, which change amino acids 219 (methionine to lysine) and 225 (cysteine to arginine) of Ycs4 (Fig. 1C).

**Figure 1.**
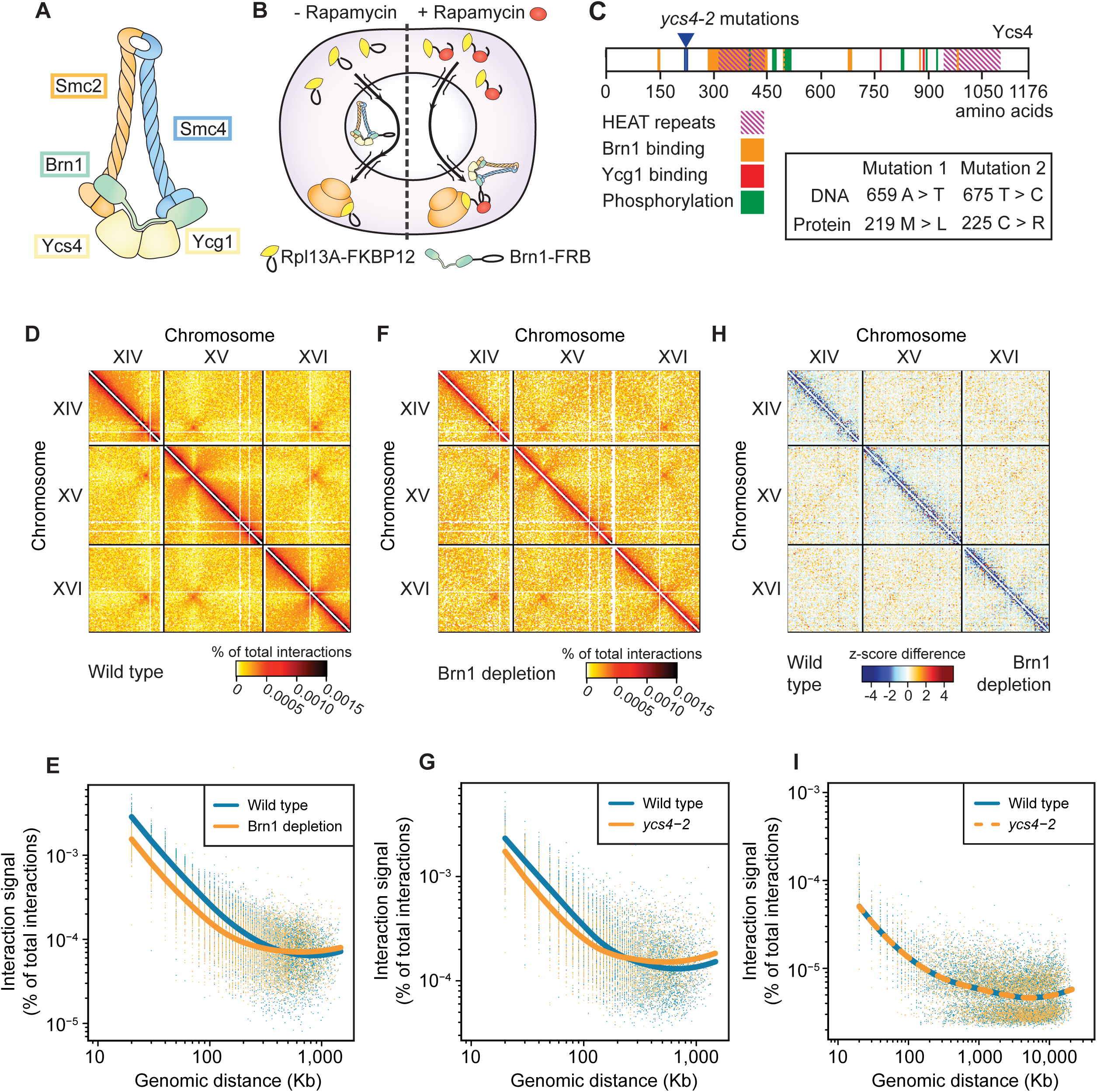
The *S. cerevisiae* genome is decompacted upon condensin perturbation. (A) Schematic of the 5 subunits that make up the condensin complex. (B) Schematic of the anchor-away approach. Upon addition of rapamycin the FRB domain fused to Brn1 binds to the FKBP12 domain fused to the ribosomal subunit Rpl13A. As a result, condensin is shuttled out of the nucleus. (C) The location of the *ycs4-2* mutations. Previously mapped structural and functional domains, and known phospho-acceptor sites are indicated. (D) Chromosome interaction frequencies across three large yeast chromosomes in wild type (parental anchor-away strain). Both wild-type and Brn1 anchor-away strains were grown asynchronously in logarithmic phase and cells were collected 30 min after rapamycin addition. (E) Decay curves plotting percentage of total interactions as a function of distance between interacting points. (F) Same as panel D, data from the Brn1 anchor-away strain. (G) Same as panel E, but from asynchronous cultures of wild-type and *ycs4-2* strains incubated two hrs at restrictive temperature. (H) Interaction difference between the wild-type and condensin-depleted cells. Z-scores for all interactions were calculated in both backgrounds. Wild-type was then subtracted from Brn1 depletion. (I) Decay curves for *C. elegans* nuclei that were spiked into the 3C-seq experiments for *ycs4-2* shown in Figure 1G and S1B.

### Shift from short to longer-range chromosomal interactions in the absence of condensing

Plotting normalized interaction frequencies between 10kb bins of the wild-type dataset showed many of the features common to chromosome conformation capture experiments in yeast and other organisms (53, 54). Interactions between loci on the same chromosome are more frequent compared to *trans* interactions (Fig 1D). Moreover, distance-decay curves show that such *cis* interaction frequencies decrease as the genomic distance between loci increases (Fig. 1E). Condensin depletion led to a reduction in local interactions with a marked increase in *trans* interactions and long-range *cis* interactions (Fig. 1F). This result is also supported by the flattened distance decay curves (Fig. 1E) and highlighted by plotting the z-score subtraction of interaction frequencies between Brn1-depleted and wild-type cells (Fig. 1H). An increase in longer-range interactions at the expense of short-range (<100 kb) was also observed in *ycs4-2* strain at the restrictive temperature (Fig. 1G, S1B).

To ensure that the change in the range of chromosomal interactions upon condensin inactivation was not due to technical differences between the wild-type and mutant samples, we used a spike-in approach. We added *C. elegans* nuclei at similar proportions to wild-type and *ycs4-2* nuclei before proceeding with the 3C-seq protocol. Reads unique to each organism were then used independently to produce decay curves of interaction frequencies within each sample. The spike-in controls showed a similar pattern of decay for interaction-frequencies in 3C-seq reactions containing wild-type and *ycs4-2* mutant nuclei (Fig. 1I), validating that condensin mutation causes a shift from short to long-range interactions in yeast.

These results are consistent with condensin promoting local compaction across the genome. The notion of global genome decompaction upon condensin inactivation in cycling cells agrees with recent depletion experiments using an auxin-dependent degron on the condensin subunit Smc2 in mitosis (37), whereas a temperature-dependent degron on Smc2 led to more limited effects (36).

### Centromere and telomeres maintain clustering in the absence of condensin

A characteristic feature of yeast genome organization is the tethered clustering of centromeres and telomeres in the nuclear periphery (55-57). These features have previously been observed through chromosome capture approaches (53, 54, 58), and were also apparent in our data. In the heat maps, the strong *trans* interactions between centromeres were observed as an array of foci (Fig. 1D). Moreover, nearly all of the 240 potential pairwise interactions between centromeres were in the top 1% of all interactions, with a significant enrichment compared to random shuffling of centromeres across the genome (Fig. 2A). Following condensin depletion, centromere clustering was maintained as previously observed with immunofluorescence (59), indicating that centromere attachment to the spindle pole is largely unaffected by condensin removal. We note, however, that condensin may have a transient role in centromere clustering at the metaphase-to-anaphase transition (37).

**Figure 2.**
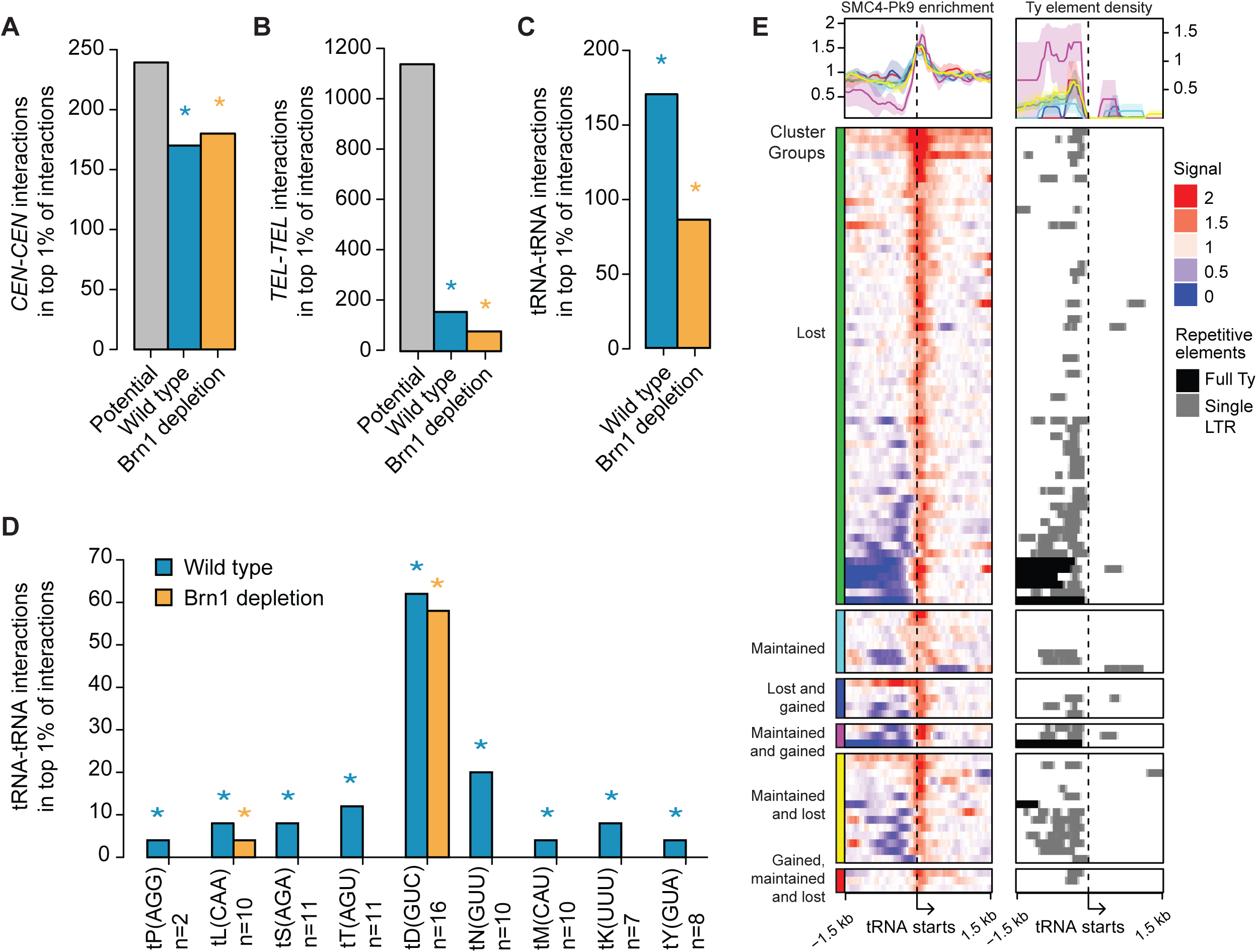
Condensin-dependent clustering of specific chromosomal loci. (A - D) Clustering of centromeres (A), telomeres (B), tRNA genes (C) and tRNA gene families (D). On the y-axis, the number of homotypic interactions within the top 1% of all interactions is plotted. ‘Potential’ shows all possible interaction pairs. Asterisks show significant enrichment of clustering compared to 1,000 permutations based on random shuffling of each set of genomic annotation coordinates (p < 0.05). (D) The tRNA gene families that show significant clustering in at least one condition are plotted (for all tRNA gene families, see Fig S2C). n = number of genes in each family. (E) (Left panel) Smc4-Pk9 ChIP-seq enrichment scores in the 3kb surrounding clustered tRNA genes (those that have interactions in the top 1% of all interactions). Lowest ChIP scores correspond to the distribution of Ty and LTR elements (right panel), possibly reflecting the low mappability to these repetitive regions. Each tRNA gene was categorized based on whether all of its interactions with other tRNA genes were lost or maintained. Some tRNA genes had a mixture of these categories and/or gained new interactions, which resulted in additional mixed categories. No tRNA genes only gained interactions, so this category is omitted.

We also observed significant enrichment of pairwise telomere interactions in wild type, though the strength of clustering was weaker compared to centromeres (Fig. 2B). Similar to centromeres, condensin depletion did not significantly alter telomere clustering in cycling cells (Fig. 2B). The condensin independence is likely due to the physical tethering of these loci to the nuclear periphery and does not reflect a general resistance of these regions to condensin-mediated effects. Accordingly, *cis* interactions of regions immediately flanking the centromeres were reduced upon condensin depletion similar to the rest of the genome (Fig. S2A, B). These data indicate that the clustering of centromeres and telomeres is refractory to the large changes in chromosomal interactions upon condensin depletion.

### Condensin is required for clustering of a subset of tRNA genes

Previous cytological analyses revealed condensin-dependent clustering and association of yeast tRNA genes with the nucleolus, the location of the rDNA (38, 60). We failed to detect significant association of tRNA genes with the rDNA (Fig. S2C, D), but tRNA-tRNA interactions were indeed enriched within the top 1% of all interactions in wild-type cells. These interactions were reduced upon condensin depletion in our data sets, suggesting a role for condensin in tRNA gene clustering (Fig. 2C).

Further analyses of homotypic clustering of tRNA genes showed differences in the extent and condensin-dependence of clustering (Fig. 2D). Of the 40 tRNA gene families, 8 exhibited significant condensin-dependent homotypic interactions (Fig. S2E), including the t^LEU^[CAA] genes used in previous FISH experiments (60). Interestingly, the largest tRNA gene family in the yeast genome (t^ASP^[GUC]) exhibited robust clustering that was unaffected by condensin inactivation (Fig. 2D). The reason for this unusual behavior is unclear because the 11 genes of this family are dispersed among other tRNA genes on multiple chromosomes. They are also not enriched near centromeres or telomeres, indicating that the observed condensin-independent clustering is not a result of hitchhiking with those features. Moreover, although tRNA genes are sites of condensin binding (25, 26, 35, 38), ChIP-seq analysis of the condensin subunit Smc4 revealed no clear correlation between relative condensin enrichment and condensin-dependent clustering (Fig. S2F), indicating that condensin binding is not sufficient to explain the specificity of tRNA gene clustering. We conclude that condensin-mediated clustering of tRNA genes is not a general phenomenon and that at least one tRNA gene family uses a condensin-independent clustering mechanism.

### Chromosome XII has a unique condensin-dependent organization

One chief point of condensin binding across the yeast genome (25, 61, 62) is the rDNA, which resides in a single ∼1Mb array in the middle of ChrXII. As observed previously, the rDNA effectively splits ChrXII into two independent interaction domains (Fig. 3A) (53, 54). Depletion of condensin led to an increase in interactions between the two ChrXII domains (Fig. 3B), whereas a strong reduction of local interactions was observed in the genomic regions abutting the rDNA (Fig. 3C). A similar result was seen in the *ycs4-2* mutant (Fig. S3A, B). These data indicate that condensin controls the conformation of the sequences flanking the rDNA array.

**Figure 3.**
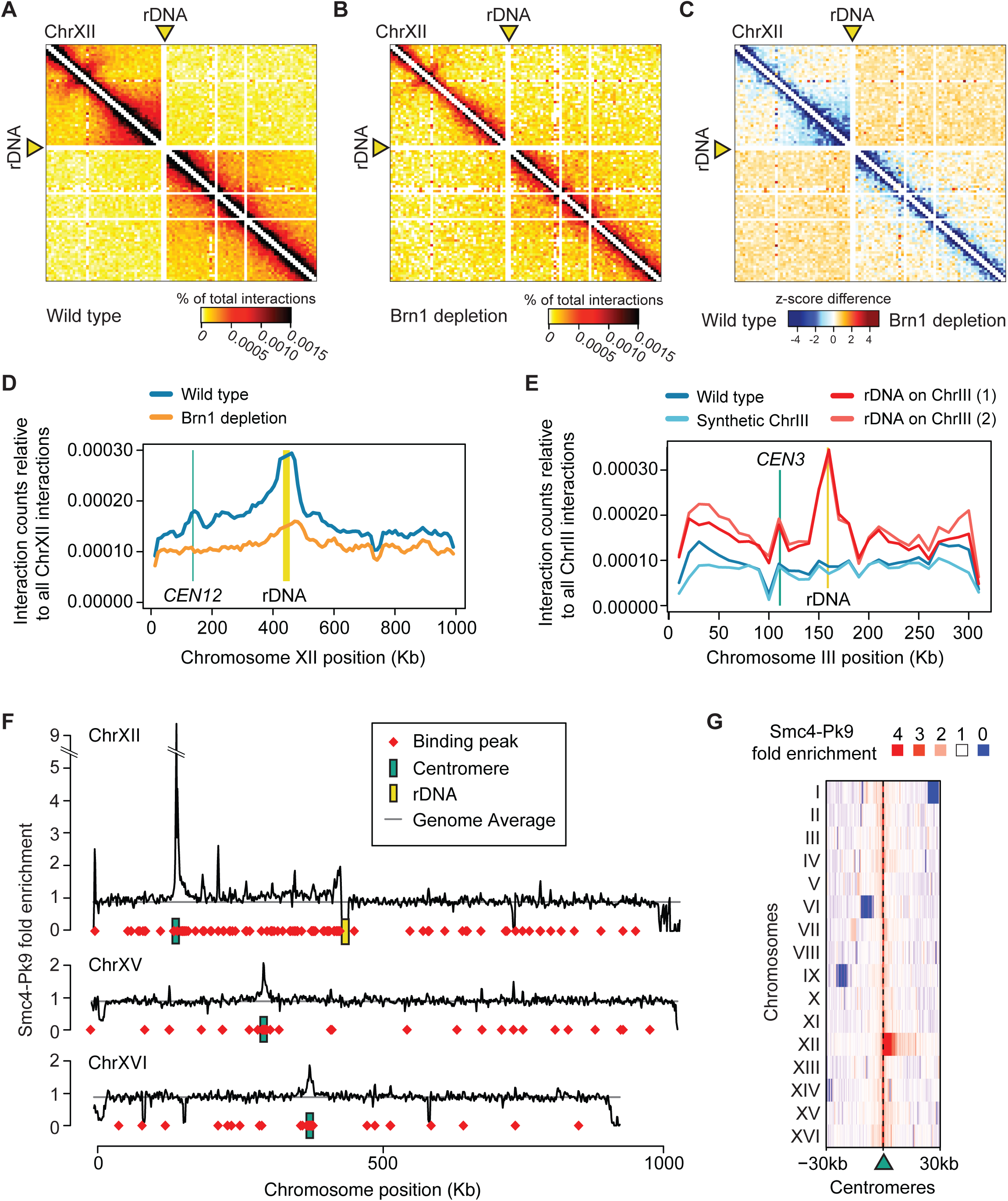
Condensin-mediated organization of the rDNA-containing chromosome XII. ChrXII interaction map from asynchronously growing yeast cells without (A) and with (B) condensin depletion using anchor-away. (C) Z-score subtraction of wild type from condensin depletion data shown in panels A and B. (D) *In silico* 4C analysis using the rDNA as bait to assess its interaction profile with the rest of ChrXII. (E) *In silico* 4C of rDNA using previously published Hi-C data (63). The red lines show profiles from two independently constructed strains in which the rDNA array was inserted into the middle of a synthetic ChrIII. The blue lines show profiles of wild-type and synthetic ChrIII. (F) Smc4-Pk9 ChIP enrichment along three similarly sized chromosomes in asynchronous cells. Data was smoothed using 30kb windows sliding over 10kb steps. Red diamonds: condensin binding peaks from MACS2 analysis; green boxes: centromeres; yellow box: rDNA. The grey line indicates genome average. (G) ChIP-seq enrichment of Smc4-Pk9 in 50bp windows across the 60kb surrounding each centromere.

Remarkably, the effect of condensin is not symmetric; the centromere-proximal left flank of the rDNA (ChrXII-L) exhibited a stronger and more far-reaching loss of local interactions upon condensin depletion than the centromere-distal right flank (ChrXII-R; Fig. S3C) (37). To test if this asymmetry is driven by polar interactions of rDNA array with either flank, we performed an *in-silico* 4C analysis. We considered the entire rDNA as a single repeat and determined its interactions with the rest of ChrXII (Fig. 3D). These analyses revealed that the rDNA interacts more with ChrXII-L than ChrXII-R in wild-type cells, with particularly strong interactions between the rDNA and *CEN12* (Fig. 3D, S3D). This contact was specific because there was no enrichment of interactions between the rDNA and the centromeres of other chromosomes (Fig. S2G). Furthermore, unlike the pairwise interactions between centromeres, the interaction between the rDNA and *CEN12* depended on condensin (Fig. 3D, S3D).

The interaction between the rDNA and *CEN12* could be the result of specialized tethering sequences near *CEN12* or of their presence on the same chromosome. To distinguish between these possibilities, we used previously published Hi-C data from a synthetic yeast strain in which the rDNA array was inserted onto ChrIII (63). Translocation of the rDNA onto ChrIII was sufficient to create two domains on ChrIII, similar to what is observed for the native ChrXII (53, 54, 63). Moreover, *in-silico* 4C analysis of this strain showed interaction between rDNA and *CEN3*, whereas the interaction with *CEN12* was lost (Fig. 3E and S3E), suggesting that the rDNA specifically interacts with the centromere of the same chromosome. It is possible that condensin-mediated tethering of rDNA to centromere (reported to occur specifically during anaphase (37)) may be important for proper segregation of the rDNA array, for which condensin function was shown to be essential (64, 65).

The condensin dependence of the rDNA-*CEN12* interaction suggested that condensin might be particularly active in the interval between these two loci. Indeed, analysis of Smc4-Pk9 binding along ChrXII revealed a specific enrichment of condensin between the rDNA and *CEN12* in wild type cells (Fig. 3F). Condensin binding is sharply delimited, as binding is at the genome-wide average on the left side of *CEN12* (Fig. 3F). No other centromere showed a comparably asymmetric condensin enrichment (Fig. 3G). These data indicate that the genomic interval between *CEN12* and the rDNA forms a domain of increased condensin binding and condensin-dependent interactions akin to TADs in higher eukaryotes.

### Condensin-dependent changes to chromosomal interactions are not coupled to global changes in gene expression

Given the broad effects of condensin on chromosome compaction, we sought to define the effect of these changes on gene expression. To quantify global changes in gene expression levels, we performed spike-in normalized mRNA-seq analysis upon condensin inactivation. Unexpectedly, there were no gross changes to global levels of gene expression following acute condensin depletion by anchor-away (Fig. 4A). Although it is possible that 30 min of condensin depletion are not sufficient to mount a robust transcriptional response, we do not favor this idea because transcriptional responses in yeast occur within minutes (66, 67) and this time frame is sufficient to observe a robust loss of compaction by 3C-seq. However, to further test this apparent resilience of the transcriptional program to genome decompaction, we also compared global expression levels in *ycs4-2* mutants after two hours of incubation at restrictive temperature. Again, no change in global gene expression was observed in response to condensin inactivation (Fig. 4B). These results suggest that the changes in chromosomal interactions observed upon condensin depletion do not lead to immediate changes in global transcription.

**Figure 4.**
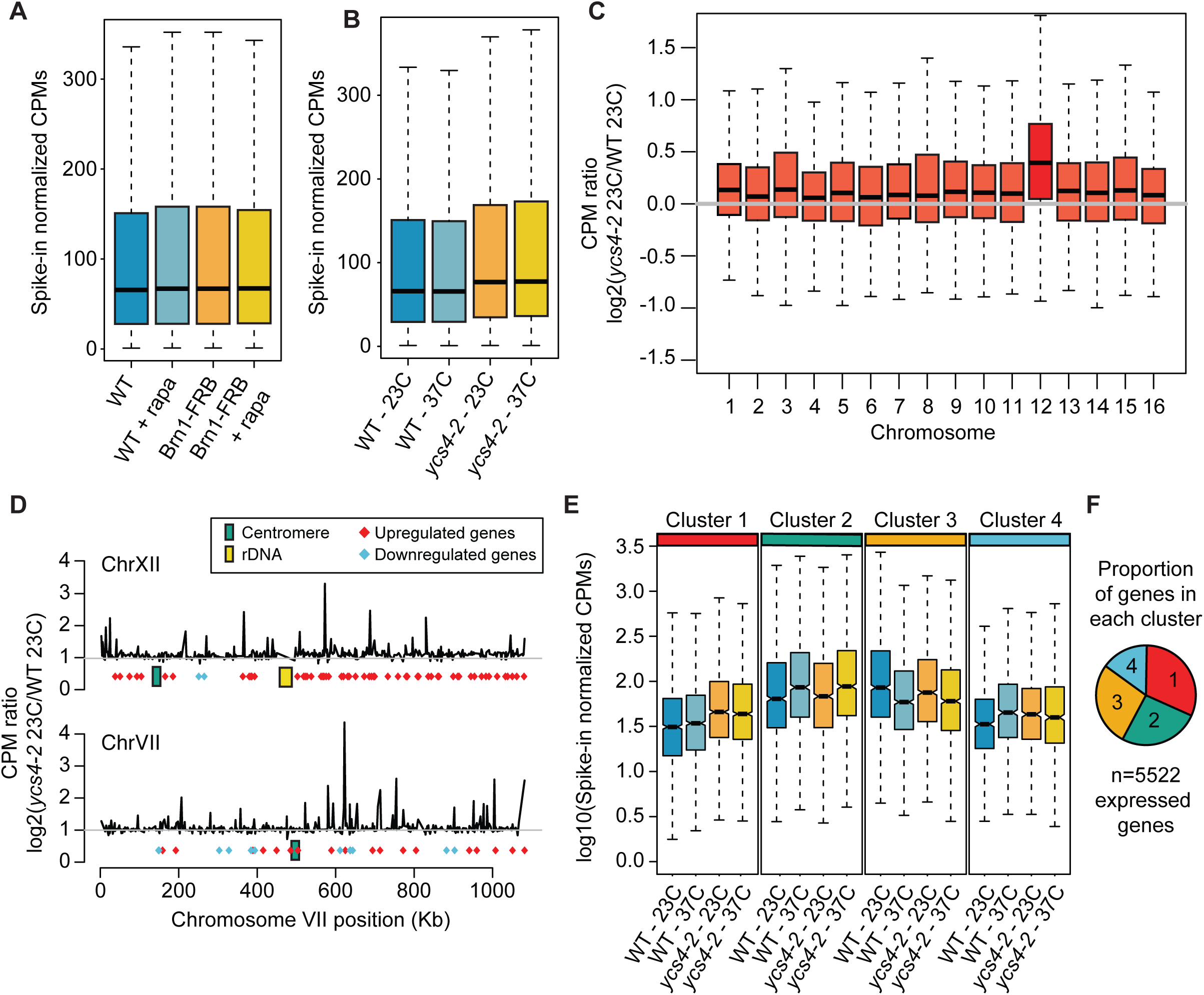
mRNA-seq analysis following condensin perturbation. (A) Spike-in corrected counts per million (CPM) for all genes are plotted. Data are from wild-type parental and Brn1 anchor-away strains grown asynchronously and collected with or without rapamycin addition of 30 min. (B) Same as panel A, but from data in the wild-type and *ycs4-2* strain incubated at permissive (23°C) or restrictive (37°C) temperature for two hours. (C) Gene expression ratios between *ycs4-2* and wild-type strain (grown at 23°C) are plotted for all chromosomes. (D) Average gene expression ratios (from panel C) within 30kb windows sliding with a 10kb step across chromosomes XII (top) and VII (bottom). Differentially expressed genes were determined by edgeR and labeled with red squares. (E) Four gene groups were generated by hierarchical clustering of gene expression from *ycs4-2* and wild-type strains grown at permissive temperature and restrictive temperature. Expression levels of genes within each cluster are shown for each growth condition. (F) Pie chart shows number of genes in each cluster.

Interestingly, the *ycs4-2* mutation was associated with a temperature-independent effect on the global level of gene expression. Both at the permissive and restrictive temperatures, the *ycs4-2* strain exhibited significantly increased average expression compared to wild type (Fig. 4B). These data suggest that the *ycs4-2* mutation is not fully functional at the permissive temperature, consistent with previous reports (34, 35). Notably, the biggest effect of *ycs4-2* on gene expression was observed on ChrXII (Fig. 4C), with a particular enrichment of upregulated genes on ChrXII-R (Fig. 4D). This spatial bias was not due to changes in copy number (Fig. S4A) or differences in the size of the rDNA array (Fig. S4B). Other possible reasons are considered in the Discussion.

To further separate the effects of temperature and the *ycs4-2* mutation, we hierarchically clustered genes across the different conditions based on expression level. This analysis revealed four major patterns of gene expression (Fig. 4E). Cluster 1 represents a substantial proportion of the genome and was sensitive to the *ycs4-2* mutation (Fig. 4E). Clusters 2 and 3 reflected temperature dependent changes to gene expression with gene expression increasing or decreasing, respectively. Lastly, cluster 4 included genes that responded to temperature in wild type but not upon *ycs4-2* mutation. Importantly, GO analysis did not find enrichment of cell-cycle controlled genes in any of the clusters, indicating that the expression changes were not secondary to a cell-cycle arrest at the restrictive temperature. Overall, almost 50% of the genome displayed increased expression in the *ycs4-2* strain compared to wild type at 23°C, the permissive temperature (Fig. 4F). Finally, we also evaluated if the four gene expression clusters capture any condensin-dependent changes in the anchor-away data. There was no difference in the expression of the four clusters upon condensin depletion by Brn1 anchor-away (Fig. S4C). We conclude that genome decompaction triggered by removal of condensin does not have immediate effects on global gene expression.

## Discussion

Here, we show that condensin is essential for the organization of the whole *S. cerevisiae* genome without controlling global gene expression levels. The primary effect of condensin depletion and mutation was a reduction in the frequency of short-range interactions, suggesting that the mechanism by which condensins mediate chromosome compaction is through promoting local interactions (<100kb). Condensin-mediated interactions at this scale are consistent with computational models of chromosome compaction before and during early prophase (68, 69). The molecular mechanism by which condensin favors short-range interactions may be based on the “loop extrusion” model, which has been gaining recent traction (70-72). In this model, a DNA loop is pushed through the condensin ring to progressively get larger. If condensin were repeatedly unloaded from chromosomes due to instability and/or slower processivity of extrusion, the result would be condensins mediating short-range interactions (69, 73). Alternatively, condensin may stabilize contacts between independent loci that occur more frequently over short distance due to the polymer nature of chromosomes (74).

Condensin also promotes specific long-range and interchromosomal interactions such as clustering of tRNA genes. A recent study in *S. cerevisiae* failed to observe the effect of condensin depletion in tRNA gene clustering (36). We predict that this result was due to insufficient condensin depletion based on the mild effect on genome-wide Hi-C interactions (36). Indeed, a second study reported greater effects of condensin depletion on Hi-C interactions in yeast (37). The notion that condensin functions at different length scales is supported by analyses in fission yeast, where condensin was shown to mediate long-range clustering of cell-cycle regulated genes in addition to local compaction (33, 75). While loop extrusion is an effective model to explain formation of chromosomal loops at different scales, how condensin mediates interchromosomal interactions such as those between certain tRNA genes remains unclear. It is possible that specific recruitment of condensins by TFIIIC and transcription factors creates condensin-mediated links between these loci (33, 38).

One possible consequence of condensin-mediated chromosomal interactions is the formation of defined TAD structures, as observed in fission yeast (33), *D. melanogaster* (76), and in *C. elegans* (24). Our data support a similar function in budding yeast, albeit at a more limited scale. In particular, condensin is required for a nested condensin-enriched domain between the rDNA and *CEN12*. Similar to TAD boundaries (77), both domain boundaries, the rDNA and *CEN12*, are highly bound by condensin. It is possible that when strong condensin recruitment domains, such as the rDNA and *CEN12*, are adjacent to each other, condensin binding is enhanced in the intervening region. Such behavior would be akin to *C. elegans* dosage compensation, where a specialized condensin complex is recruited to specific recruitment sites, from which it spreads out (78).

In *C. elegans* and other organisms, condensins have been linked to the regulation of transcription (38, 79, 80). Our data indicate that the global gene expression program is largely unaffected by condensin depletion in yeast. We did observe an overall increase in gene expression in the *ycs4-2* strain but these effects were independent of temperature and thus not linked to increased chromosome decompaction upon condensin inactivation. As the *ycs4-2* mutation already causes transcriptional phenotypes at room temperature (34, 35), these differences could be an indirect result of the chronic presence of the condensin mutation as opposed to the acute response to rapid depletion of condensin from the nucleus by the anchor-away system. Furthermore, the *ycs4-2* strain may have adapted transcriptionally to compensate the effect of the condensin mutation. Indeed, the *YCS4* gene is located on ChrXII-R, within the region containing an enrichment of genes that were upregulated in the *ycs4-2* strain, and mRNA levels of the *ycs4-2* allele were elevated approximately 2-fold compared to wild type.

The observation that acute depletion of condensin caused immediate effects on genome organization but not on gene expression suggests that transcription can be decoupled from global chromosome structure. We note, however, that chromosome condensation in *S. cerevisiae* is substantially less dramatic than observed for mitotic chromosomes in larger eukaryotes (81). Furthermore, gene regulatory elements in yeast are localized near the promoters and may not depend on specific chromosomal interactions as enhancers and promoters in bigger eukaryotic genomes. Finally, other energy-dependent chromatin organizers, including other SMC proteins may compensate for the loss of condensin and provide robustness against acute changes in condensin activity.

## Acknowledgements

We thank H Yu for sharing strains, M Imakaev for help establishing the analysis pipeline, D Corrales and H He for help with *S. pombe* culture and primers, and the NYU Department of Biology Sequencing Core for technical assistance and data processing. This work was supported by a fellowship from Boehringer Ingelheim Fonds to MRP, and NIH grants R01GM111715 to AH and R01GM107293 to SE.

